# Physiological characterization of nitrate ammonifying bacteria isolated from rice paddy soil via a newly developed high-throughput screening method

**DOI:** 10.1101/2020.05.06.081935

**Authors:** Hokwan Heo, Miye Kwon, Bongkeun Song, Sukhwan Yoon

## Abstract

Dissimilatory nitrate/nitrite reduction to ammonium (DNRA) has recently gained attention as a nitrogen retention pathway that may potentially be harnessed to alleviate nitrogen loss resulting from denitrification. Until recently, ecophysiology of DNRA bacteria inhabiting agricultural soils has remained largely unexplored, due to the difficulty in targeted enrichment and isolation of DNRA microorganisms. In this study, >100 microbial isolates capable of DNRA have been isolated from rice paddy soil with apparent dominance of denitrification using a novel high-throughput screening method. Six of these isolates, each assigned to a disparate genus, was examined to improve understanding of DNRA physiology. All isolates carried *nrfA* and/or *nirB*, and an isolate affiliated to *Bacillus* possessed a clade II *nosZ* gene and was capable of N_2_O reduction. A common prominent physiological feature observed in all DNRA isolates was NO_2_^−^ accumulation observed before NH_4_^+^ production, which was further examined with *Citrobacter* sp. DNRA3 (possessing *nrfA* and *nirB)* and *Enterobacter* sp. DNRA5 (possessing only *nirB*). In both organisms, NO_2_^−^-to-NH_4_^+^ reduction was inhibited by submillimolar NO_3_^−^, and *nrfA* or *nirB* transcription was down-regulated when NO_3_^−^ was being reduced to NO_2_^−^. Both batch and chemostat incubations of these isolates with excess organic electron donors produced NH_4_^+^ from reduction of NO_3_^−^; however, incubation with excess NO_3_^−^ resulted in NO_2_^−^ buildup but no substantial NH_4_^+^ production, presumably due to NO_3_^−^ presence. This previously overlooked link between NO_3_^−^ repression of NO_2_^−^-to-NH_4_^+^ reduction and the C-to-N ratio regulation of DNRA activity may be a key mechanism underpinning denitrification-vs-DNRA competition in soil.

**IMPORTANCE:** Dissimilatory nitrate/nitrite reduction to ammonium (DNRA) is an anaerobic microbial pathway that competes with denitrification for common substrates NO_3_^−^ and NO_2_^−^. Unlike denitrification leading to nitrogen loss and N_2_O emission, DNRA reduces NO_3_^−^ and NO_2_^−^ to NH_4_^+^, a reactive nitrogen with higher tendency to be retained in soil matrix. Therefore, stimulation of DNRA has often been proposed as a strategy to improve fertilizer efficiency and reduce greenhouse gas emissions. Such attempts have been hampered by lack of insights into soil DNRA ecophysiology. Here, we have developed a novel high-throughput screening method for isolating DNRA-catalyzing organisms from agricultural soils without apparent DNRA activity. Physiological characteristics of six DNRA isolates were closely examined, disclosing a previously overlooked link between NO_3_^−^ repression of NO_2_^−^-to-NH_4_^+^ reduction and the C-to-N ratio regulation of DNRA activity, which may be key to understanding why significant DNRA activity is rarely observed in nitrogen-rich agricultural soils.

## INTRODUCTION

Nitrogen is an essential element for plant growth. Today, the Haber-Bosch process, used primarily for production of nitrogen fertilizers, is singled out as the largest energy-consuming industrial process, with global energy consumption summing up to 2.5% of the total energy consumed across the globe, and naturally, is one of the largest sources of greenhouse gases (1, 2). The increased nitrogen flux in the soil and aquatic environments, as a consequence of fertilizer application to agricultural soils, has also led to aggravation of various nitrogen-related environmental problems, e.g., enrichment of NO_3_^−^ in groundwater and harmful algal blooms as a symptom of eutrophication in surface water (3). Thus, mitigation of the ‘nitrogen dilemma’ has been regarded as one of the most impending issues for environmental sustainability (4).

Despite the environmental consequences, nitrogen is not used efficiently in agroecosystems. Nitrogen fertilizer efficiency, i.e., the proportion of applied fertilizer nitrogen that eventually ends up in crop biomass rarely exceeds 40%, due to nitrogen loss via sequential nitrification and denitrification reactions (5). These microbial reactions are also the major culprits of N_2_O emissions. Several different strategies have been devised to limit nitrogen loss and N_2_O emissions from soil systems, including the use of nitrification inhibitors and slow-release fertilizers (6, 7). Another possible strategy recently proposed for improved soil nitrogen management is to outcompete the denitrification pathway with nitrogen-retaining DNRA (dissimilatory nitrate/nitrite reduction to ammonium) pathway (8-11). The DNRA pathway is the reduction of NO_3_^−^/NO_2_^−^ to NH_4_^+^ catalyzed by the microorganisms equipped with cytochrome *c*_552_ nitrite reductases (encoded by *nrfA* genes) or NADH-dependent nitrite reductases (encoded by *nirB* genes) generally perceived as assimilatory nitrite reductases (12). According to current limited knowledge, NO_2_^−^-to-NH_4_^+^ reduction may serve as the electron acceptor reaction for respiration (respiratory DNRA) or electron dump for NADH regeneration in fermentation of complex organics (fermentative DNRA) (13-15).

Denitrification and DNRA pathways compete for common substrates, NO_3_^−^/NO_2_^−^, and thus, stimulating one would repress the other (16, 17). Previous investigations suggested that DNRA is favored in environments with high organic carbon (C) content and limiting supply of nitrogenous electron acceptors (NO_3_^−^/NO_2_^−^) (18-20). This hypothesis was further corroborated by recent laboratory experiments with microbial enrichments and axenic microbial cultures harboring both denitrification and DNRA pathways; however, conflicting observations (e.g., in experiments with *Intrasporangium calvum* and the *Deltaproteobacteria-dominated* wastewater enrichments) suggest the possibility that the observed correlation between DNRA activity and the C-to-N ratio (the ratio of C in bioavailable organic compounds to N in NO_3_^−^/NO_2_^−^ in this context) may be circumstantial (17, 21-23).

Microbial population responsible for DNRA in soils can now be analyzed without culturing, using recently developed molecular tools targeting *nrfA* genes or metagenomics and meta-omics techniques (24, 25). Nevertheless, these culture-independent analyses need to be complemented with culture-based examinations into physiology of soil DNRA isolates, especially so considering that the previous observations that *nrfA* abundance may be decoupled from DNRA activity and that *nirB*-possessing organisms may also contribute to the overall DNRA activity (26, 27). The difficulty in securing DNRA isolates from soils with apparent dominance of denitrification has been the bottleneck in culture-based investigation of DNRA physiology. The traditional approach that has been used for isolation of DNRA bacteria was NO_3_^−^-amended anoxic enrichment followed by isolation via serial dilution and/or single colony picking and screening for isolates capable of NH_4_^+^ production from anaerobic incubation with NO_3_^−^. This approach would require extensive menial labor for screening out a few DNRA isolates from overwhelming number of denitrifiers when applied to soils with apparent dominance of denitrification. In fact, in the only reported case of such targeted isolation of soil DNRA organisms, only three out of 80 NO_3_^−^-reducing isolates were revealed as capable of reducing NO_3_^−^ to NH_4_^+^ (28). Not surprisingly, the model organisms examined for DNRA ecophysiology rarely originated from soil (15, 17, 29-32). Further, most of these isolates had been aerobically isolated and cultured for decades in laboratory settings before they were recognized as being capable of DNRA. Thus, use of these isolates as representatives of soil DNRA bacteria have received criticism as lacking ecological relevance to the fate of NO_3_^−^ in anoxic agricultural soils.

To address this ecological relevance issue in examining soil DNRA ecophysiology, a less onerous and time-consuming method for isolation of DNRA bacteria in denitrification-dominant agricultural soils was in need. Here, to address this issue,a rapid inexpensive high-throughput screening method was developed utilizing the well-established salicylate method for NH_4_^+^ detection and quantification (33). Reductive transformation of NO_3_^−^ was examined with six DNRA organisms isolated from rice paddy soils using this novel screening method, affiliated to *Bacillus* (belonging to *Firmicutes* phylum), *Aeromonas, Citrobacter, Enterobacter, Klebsiella* and *Shewanella* (belonging to *Proteobacteria* phylum) genera. Their most obvious common physiological feature was NO_3_^−^ inhibition of NO_2_^−^-to-NH_4_^+^ reduction, which had been also previously observed with *nrfA*-and-*nirB*-harboring organisms *Escherichia coli* and *Bacillus vireti* (34, 35). With a series of batch and continuous culture experiments, we identified this NO_3_^−^ repression of DNRA activity as one of the mechanisms underpinning the widely acknowledged but controversial C-to-N ratio regulation of DNRA-vs-denitrification competition.

## RESULTS

### Isolation of DNRA bacteria from denitrification-dominant agricultural soil

Out of 192 colonies each from the lactate- and glucose-amended rice paddy soil enrichments, both with negligible NH_4_^+^ production from NO_3_^−^ reduction, 126 and 12 colonies tested DNRA-positive (Fig. S1). Sequencing of the 16S rRNA gene amplicons of the positive colonies (30 and 12 colonies from lactate- and glucose-amended enrichments, respectively) identified six bacterial genera: *Aeromonas*, *Bacillus*, *Shewanella* (lactate-amended), *Enterobacter*, *Klebsiella* (glucose-amended), and *Citrobacter* (both) (Fig. S2). The DNRA activities in six of these isolates, each belonging to a different genus, were further examined.

### Identification of functional genes relevant to dissimilatory nitrogen reduction

The draft genomes of the six DNRA isolates were constructed from the HiSeq sequencing reads (the sequencing statistics presented in Table S2). The functional genes potentially relevant to turnover of reactive nitrogen species or regulation of nitrogen metabolism were then analyzed in these draft genomes (Fig. 1 and Table S3). The isolates that originated from lactate-enriched cultures all possessed *nrfA* genes encoding NH_4_^+^-forming cytochrome *c*_552_ nitrite reductases, while the two isolates from glucose-enriched cultures lacked *nrfA* gene, but possessed *nirB* genes, suggesting that NirB-type nitrite reductase was responsible for dissimilatory reduction of NO_2_^−^ to NH_4_^+^ in these organisms. Only *Citrobacter* sp. DNRA3 possessed both *nrfA* and *nirB*. All six isolates have *napA* in their genomes, and *Citrobacter* sp. DNRA3*, Enterobacter* sp. DNRA5, and *Klebsiella* sp. DNRA6 carry *narG*, indicating the genomic potential of these organisms to reduce NO_3_^−^ to NO_2_^−^. Neither *nirK* nor *nirS*, i.e., genes encoding NO-forming nitrite reductases, was present in any of the isolates; however, a clade II *nosZ* gene was identified in the draft genome of *Bacillus* sp. DNRA2, suggesting N_2_O-reducing capability.

**Fig. 1.**
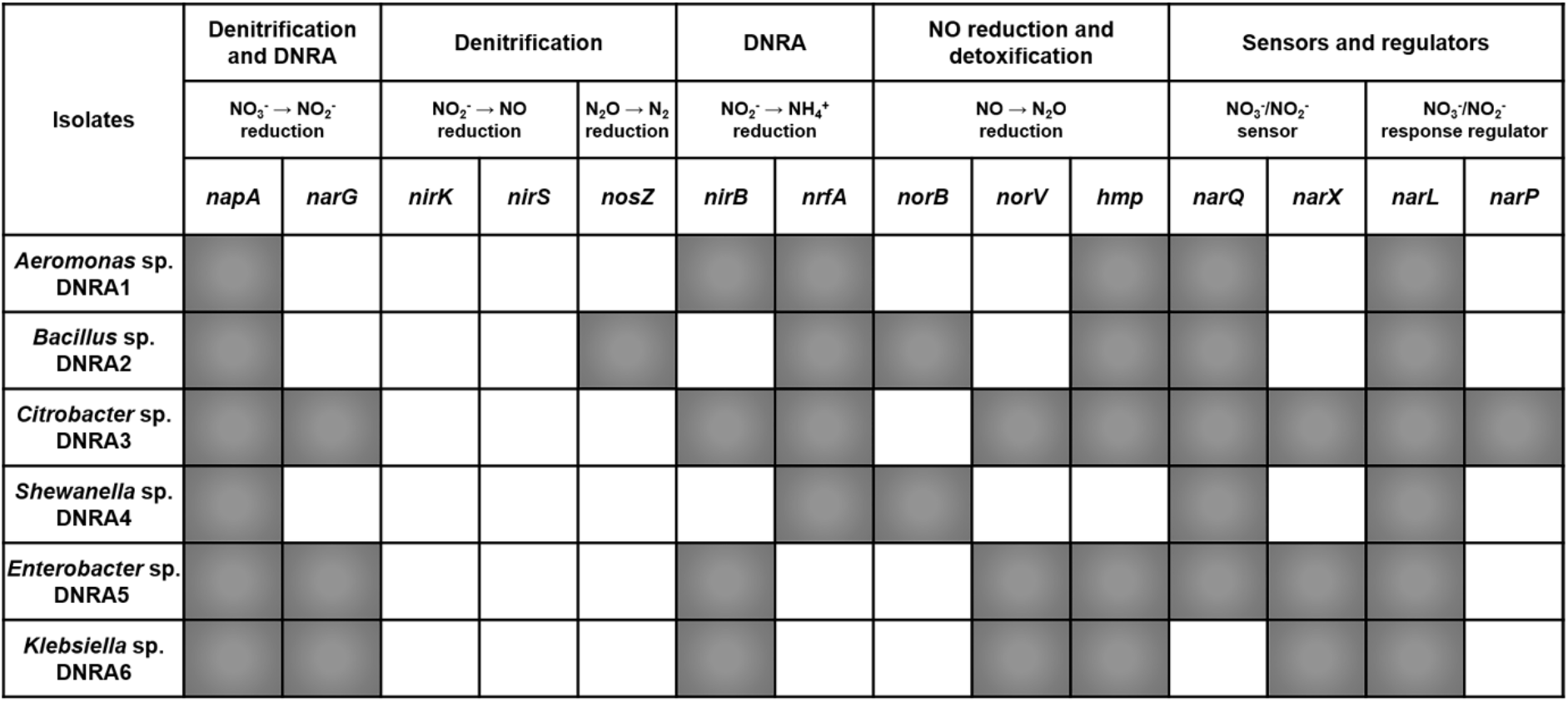
Functional genes identified in the draft genomes of the six DNRA bacteria that are potentially relevant to turnover of reactive nitrogen species or regulation of dissilatory nitrogen metabolism. The genes that were recovered in the draft genome are represented as shaded boxes.

### NO_3_^−^ reduction of the candidate DNRA isolates

Reductive transformation of NO_3_^−^ was observed with the axenic cultures of the six candidate DNRA isolates with or without 10% C_2_H_2_ in the headspace (Fig. 2, S3, and S4). All six isolates completely reduced the initially supplemented NO_3_^−^ to NK’ via NO_2_^−^ with lactate or glucose as the source of electrons. Lactate-coupled NO_3_^−^ reduction in *Aeromonas* sp. DNRA1, *Bacillus* sp. DNRA2, *Citrobacter* sp. DNRA3, and *Shewanella* sp. DNRA4 resulted in near-stoichiometric production of NH_4_^+^ from NO_3_^−^. Reduction of NO_3_^−^ to NH_4_^+^ was also observed in *Enterobacter* sp. DNRA5 and *Klebsiella* sp. DNRA6 grown on glucose; however, produced NH_4_^+^ only amounted to 36.3±1.1 and 32.1±0.2 μmoles, respectively, which were less than half of nitrogen added as NO_3_^−^. As the cell density of the glucose-fed *Enterobacter* sp. DNRA5 and *Klebsiella* sp. DNRA6 reached at least 2.5-fold higher than the lactate-consuming isolates, the missing nitrogen was likely due to assimilation. Despite the absence of *nirK* or *nirS* gene, N_2_O production was observed in all of the examined isolates during NO_3_^−^ reduction when incubated with C_2_H_2_. The amounts of N_2_O produced varied across the isolates, ranging from 0.40±0.06 μmoles N_2_O-N of *Shewanella* sp. DNRA4 to 3.5±0.3 μmoles N_2_O-N of *Citrobacter* sp. DNRA3. In all six isolates examined, the start of N_2_O production corresponded with the start of NH_4_^+^ production, suggesting that N_2_O was the byproduct of NO_2_^−^-to-NH_4_^+^ reduction, not NO_3_^−^-to-NO_2_^−^ reduction. Of the six isolates, only *Bacillus* sp. DNRA2 showed a substantially different N_2_O-N time series profile when incubated without C_2_H_2_ (Fig. S4). The absence of N_2_O accumulation suggested that N_2_O consumption occurred simultaneously with DNRA in this clade II *nosZ*-harboring organism.

**Fig. 2.**
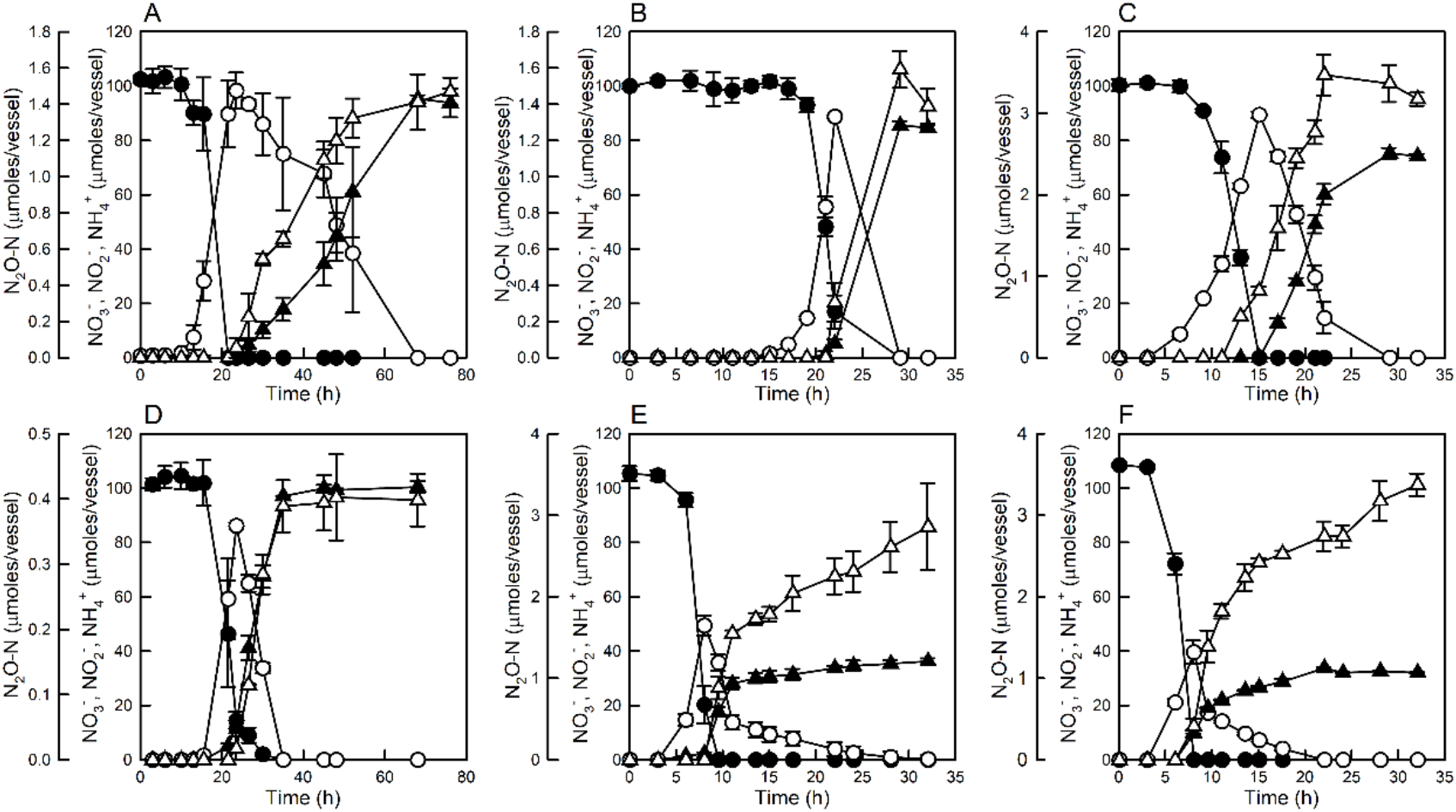
NO_3_^−^ reduction monitored in 100-mL batch cultures (prepared in sealed 160-mL serum bottles with headspace consisting of 90% N_2_ and 10% C_2_H_2_) of (A) *Aeromonas* sp. DNRA1, (B) *Bacillus* sp. DNRA2, (C) *Citrobacter* sp. DNRA3, (D) *Shewanella* sp. DNRA4, (E) *Enterobacter* sp. DNRA5, and (F) *Klebsiella* sp. DNRA6. The averages of biological replicates (*n*=3) are presented with the error bars representing their standard deviations (NO_3_^−^: ●, NO_2_^−^: ○, NH_4_^+^: ▲, N_2_O-N: Δ)

Accumulation of NO_2_^−^ before reduction to NH_4_^+^was consistently observed in all six of these isolates. Reduction of NO_2_^−^ to NH_4_^+^ did not commence until >80% of NO_3_^−^ was consumed in any of the six isolates, suggesting that NrfA- or NirB-catalyzed NO_2_^−^-to-NH_4_^+^ reduction was affected by changing NO_2_^−^ or NO_3_^−^ concentrations. The DNRA reactions of *Citrobacter* sp. DNRA3 and *Enterobacter* sp. DNRA5 were further investigated to identify whether possible causality exists between NO_2_^−^ or NO_3_^−^ concentration and DNRA activity (Fig. 3A-B). Transcription levels of *nrfA* in *Citrobacter* sp. DNRA3 and *nirB* in *Enterobacter* sp. DNRA5 were significantly higher (*p*<0.05) after NO_3_^−^ was depleted than before. Transcription of *nrfA* in *Citrobacter* sp. DNRA3 increased significantly (*p*<0.05) from 1.0±0.6 *nrfA/recA* at t=9 h (0.16±0.03 mM NO_3_^−^ and 0.82±0.03 mM NO_2_^−^ remaining) to 6.3±0.2 *nrfA/recA* at t=15 h (0.33±0.08 mM NO_2_^−^ remaining). No significant change was observed with *nirB* transcription (2.04±0.19 *nirB/recA* at t= 9 h and 1.61±0.37 *nirB/recA* at t=15 h), suggesting that NirB-type nitrite reductase was irrelevant to respiratory DNRA. Transcription of *nirB* in *Enterobacter* sp. DNRA5 followed a similar trend with that of *nrfA* in *Citrobacter* sp. DNRA3, increasing significantly after depletion of NO_3_^−^ (*p*<0.05). Substrate (NO_2_^−^) regulation of transcription was unlikely for either *nrfA* in *Citrobacter* sp. DNRA3 and *nirB* in *Enterobacter* sp. DNRA5, as the transcriptions of these genes appeared unresponsive to elevated NO_2_^−^ concentrations, as long as NO_3_^−^ was present in the medium at >0.15 mM. Thus, NO_3_^−^ concentration was the most probable environmental factor that affected the transcription of the genes encoding these DNRA-catalyzing nitrite reductases. The significant differences in the rates of NO_2_^−^ reduction measured with *Citrobacter* sp. DNRA3 or *Enterobacter* sp. DNRA5 cells harvested before and after the NO_3_^−^ depletion and treated with chloramphenicol also supported that expression of the NHAforming nitrite reductases were down-regulated by the presence of NO_3_^−^ (Fig. 3C-F). *Citrobacter* sp. DNRA3 cells extracted before NO_3_^−^ depletion did not exhibit significant NO_2_^−^ reduction activity, while the cells extracted after NO_3_^−^ depletion readily reduced NO_2_^−^ at a rate of 170±13 μmoles s^−1^ mg protein^−1^. NO_2_^−^ reduction by *Enterobacter* sp. DNRA5 cells was also ~6 times higher with the cells harvested after NO_3_^−^ depletion (101±10 μmoles s^−1^ mg protein^−1^) than the cells harvested before NO_3_^−^ depletion (17.2±5.7 μmoles s^−1^ mg protein^−1^; *p*<0.05).

**Fig. 3.**
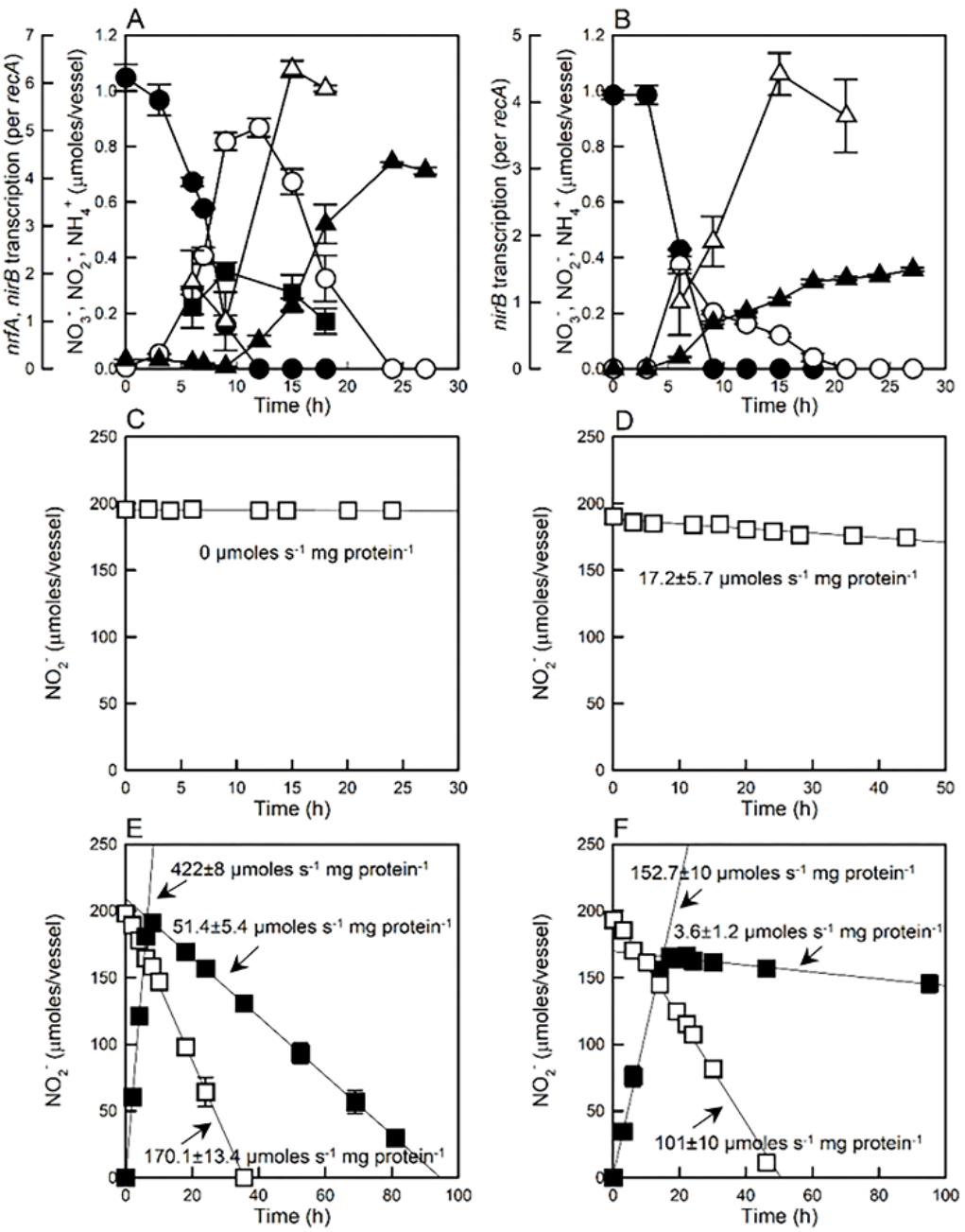
Transcription of (A) *nrfA* and *nirB* in *Citrobacter* sp. DNRA2 and (B) *nirB* in *Enterobacter* sp. DNRA5 cells as 1 mM NO_3_^−^ (●) was reduced to NH_4_^+^ (▲) via NO_2_^−^ (○). Changes to the amounts of NO_2_^−^ were monitored in the chloramphenicol-treated resting cultures of *Citrobacter* sp. DNRA2 (C, E) and *Enterobacter* sp. DNRA5 (D, F) harvested before (C, D) and after (E, F) NO_3_^−^ depletion and resuspended in fresh medium containing 2 mM NO_2_^−^ (□) or 2 mM NO_3_^−^ (■). All experiments were performed in biological replicates (*n*=3) and the error bars represent the standard deviations

In the resting-cell experiments with 2 mM NO_3_^−^ added to chloramphenicol-treated *Citrobacter* sp. DNRA3 cells harvested after NO_3_^−^ depletion, NO_2_^−^ accumulated up to 1.93±0.04 mM at a rate of 422±8 μmoles s^−1^ mg protein^−1^ before it was consumed at a rate of 51.4±5.3 μmoles s^−1^ mg protein^−1^ (Fig. 3E). The negligible NO_2_^−^ reduction activity before NO_3_^−^ depletion suggested an additional NO_3_^−^-mediated inhibitory mechanism on NrfA-type nitrite reductase activity apart from transcriptional regulation of *nrfA* gene. Such repression of NO_2_^−^ reduction activity by NO_3_^−^ presence was not observed in the parallel experiment performed with *Enterobacter* sp. DNRA5 lacking *nrfA* and presumably utilizing NirB-type nitrite reductase (Fig. 3F).

### DNRA reaction at varying C-to-N ratios: batch and continuous cultivation

*Citrobacter* sp. DNRA3 and *Enterobacter* sp. DNRA5 were grown in batch and continuous cultures, each with two different C-to-N ratios, and NO_3_^−^ reduction was monitored to investigate whether the generally perceived positive correlation between C-to-N ratio and DNRA activity may be related to the NO_3_^−^-repression of NO_2_^−^-to-NH_4_^+^ reduction (Fig. 4 and 5). When grown at the initial C-to-N ratio of 75 in batch cultures, *Citrobacter* sp. DNRA3 produced NH_4_^+^ from NO_3_^−^ reduction, and each addition of 20 μmoles NO_3_^−^ resulted in near-stoichiometric turnover to NH_4_^+^. Contrastingly, *Citrobacter* sp. DNRA3 grown at the initial C-to-N ratio of 0.3 did not result in significant increase in NH_4_^+^ concentration, but led to stoichiometric NO_2_^−^ accumulation, as NO_3_^−^ reduction produced 0.87±0.06 mM NO_2_^−^ when the initial reaction stopped at t=16 h due to depletion of lactate. Reduction of NO_3_^−^ to NO_2_^−^ resumed after replenishment with 0.2 mM lactate at t=21 h. At this low C-to-N ratio incubation condition, NO_3_^−^ was present in the culture medium throughout incubation, and the presence of NO_3_^−^ was likely the reason for the absence of *sensu stricto* DNRA activity.

**Fig. 4.**
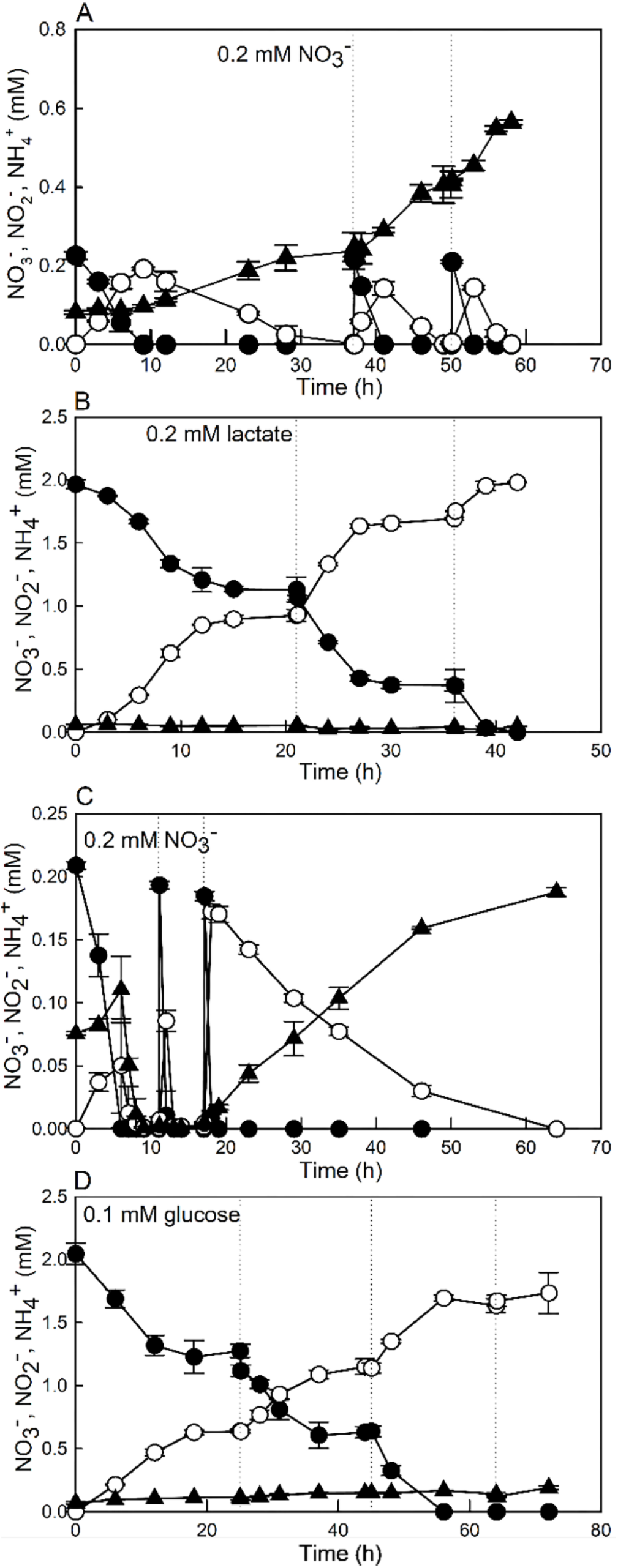
NO_3_^−^ reduction observed with batch cultures of *Citrobacter* sp. DNRA2 (A, B) and *Enterobacter* sp. DNRA5 (C, D) prepared with two different initial C-to-N ratios. The high C-to-N conditions were prepared with 0.2 mM NO_3_^−^ and 5 mM lactate (A) or 2.5 mM glucose (C), and NO_3_^−^was replenished to 0.2 mM upon NO_3_/NO_2_^−^ depletion. The low C-to-N conditions were prepared with 2.0 mM NO_3_^−^ and 0.2 mM lactate (B) or 0.1 mM glucose (D) and the carbon sources were replenished when NO_3_^−^/NO_2_^−^ reduction stopped. The dotted lines denote the time points where the limiting nutrients were replenished. The averages of biological replicates (*n*=3) are presented with the error bars representing their standard deviations (NO_3_^−^: ●, NO_2_^−^: ○, NH_4_^+^: ▲, N_2_O-N: **Δ**)

**Fig. 5.**
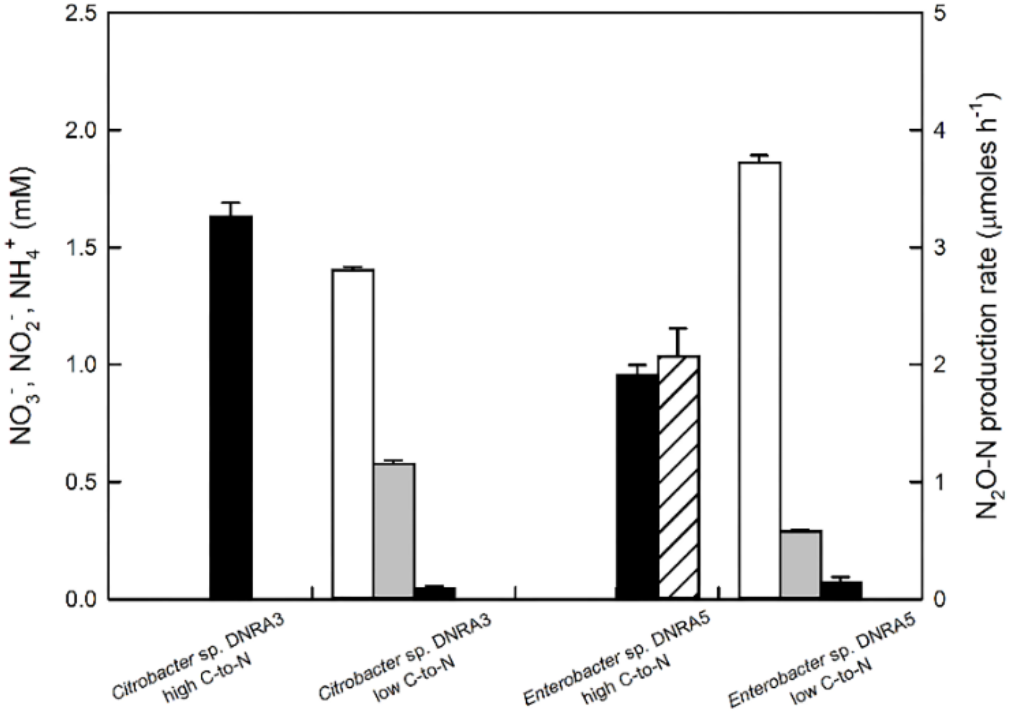
Steady-state concentrations of NO_3_^−^ (white), NO_2_^−^ (gray), and NH_4_^+^ (black) in the electron acceptor-limiting (high C-to-N ratio) and electron donor-limiting (low C-to-N ratio) chemostat cultures of *Citrobacter* sp. DNRA2 and *Enterobacter* sp. DNRA5. N_2_O-N production rate (hatched) is also presented for *Enterobacter* sp. DNRA5 cultivated under the electron acceptor-limiting condition, which was the only reactor culture with observed N_2_O production. The average values of the three measurements taken with six-hour intervals are presented with the error bars represent their standard deviations

*Enterobacter* sp. DNRA5, incubated on glucose at the C-to-N ratio of 75 produced significant amount of NH_4_^+^ only after initially added glucose (2.19±0.8 mM) was fully consumed, suggesting that substantial portions of NO_3_^−^ and its reduction products, NO_2_^−^ and NH_4_^+^, were assimilated. Upon third addition of 0.2 mM NO_3_^−^, with no glucose remaining in the medium, the sequential NO_3_^−^-to-NO_2_^−^-to-NH_4_^+^ reduction was stoichiometric, suggesting that NirB-catalyzed NO_2_^−^-to-NH_4_^+^ reduction was coupled to oxidation of the fermentation products. At the low C-to-N ratio, where the culture medium was replenished with 0.1 mM glucose upon halt in NO_3_^−^ reduction, the time series profiles of the N-species concentrations were indistinguishable from that of *Citrobacter* sp. DNRA3, save for the imperfect stoichiometry between consumed NO_3_^−^ and produced NO_2_^−^ and the modest, albeit significant, increase in NH_4_^+^ concentration from 0.07±0.01 mM at t=0 h to 0.19±0.01 mM at t=72 h. The modest production of NH_4_^+^ was in line with the reduced, but still significant NO_2_^−^ reduction rate observed in the resting-cell cultures of *Enterobacter* sp. DNRA5 extracted before NO_3_^−^ depletion.

The continuous culture of *Citrobacter* sp. DNRA3 fed medium carrying 10 mM lactate and 2 mM NO_3_^−^ (C-to-N ratio of 15), after attaining steady state, contained 1.68±0.09 mM NH_4_^+^ as the only dissolved inorganic nitrogen, indicating that NO_3_^−^ and NO_2_^−^ were readily reduced in the chemostat. When the reactor was fed the medium carrying 0.2 mM lactate and 2 mM NO_3_^−^ (C-to-N ratio of 0.3), 1.41±0.04 mM NO_3_^−^ remained in the medium at steady state, due to carbon limitation. That NO_2_^−^ was the major product of NO_3_^−^ reduction (0.58±0.04 mM at steady state) and NH_4_^+^ concentration did not significantly differ from the concentration in the fresh medium (p>0.05) indicated DNRA did not proceed further than NO_2_^−^. Similarly, with *Enterobacter* sp. DNRA5, significant NH_4_^+^ formation was observed only in the continuous culture operated at the electron-acceptor-limiting condition, i.e., at C-to-N ratio of 15. The steady state NH_4_^+^ concentration was 0.95±0.04 mM in this chemostat. In the continuous culture operated at electron-donor-limiting condition, NO_2_^−^ was the only dissolved nitrogen species with significantly higher concentration than the influent medium. Production of N_2_O (2.07±0.24 μmoles hr^−1^) was observed only in the high C-to-N chemostat of *Enterobacter* sp. DNRA5. The absence of significant NO_2_^−^-to-NH_4_^+^ reduction at the low C-to-N-ratio batch and chemostat cultures, regardless mediated by NrfA-type or NirB-type nitrite reductase, could be best explained as the inhibitory effect of NO_3_^−^.

## DISCUSSION

For long, DNRA has been recognized as one of the key reactions determining the fate of reactive nitrogen in the environment (36-38). Recovery of ^15^NH_4_^+^ from ^15^NO_3_^−^ reduction in both *in situ* column studies and *ex situ* soil incubation experiments supported that the DNRA pathway is a significant dissimilatory reaction pathway in soil environments; however, fertilized agricultural soils with the DNRA pathway dominating over the denitrification pathway, i.e., with higher NH_4_^+^ production than N_2_O+N_2_ production from NO_3_/NO_2_^−^ reduction, have rarely been reported (8, 16, 39). Thus, few attempts have been made to isolate and examine DNRA organisms from agricultural soils, and interpretations of DNRA in ecological contexts, particularly with regards to the fate of fertilizer N, have relied on extrapolation of findings from experiments with limited number of isolates, most, if not all, acquired from non-soil environments (15, 17, 28, 32, 34, 40). The high-throughput DNRA screening method developed in this study enabled easy and rapid targeted isolation by systematically combining widely-used culturing and colorimetric detection techniques. The most conspicuous benefit of the method is the ease of isolating the DNRA organisms that had been considered to be difficult to enrich from soils with apparent domination of denitrification over DNRA. Examination of the physiology of the indigenous DNRA organisms in denitrification-dominant soils, isolated using this targeted isolation technique, would be a sensible approach to investigate the reasons why NH_4_^+^ production via DNRA is often underrepresented in the nitrogen cycling of agricultural soils. Such knowledge would be immensely useful in developing soil management techniques for enhancing the nitrogen-retaining DNRA pathway (9, 39).

The newly acquired soil DNRA isolates were assigned to six genera according to their 16S rRNA gene sequences. Several of these genera have been previously confirmed to include strains capable of carrying out DNRA (*Bacillus*, *Citrobacter*, *Enterobacter*, and *Klebsiella*). The list also included a genus that has not yet been previously isolated from soil environments (*Shewanella*) and a genus without physiologically confirmed DNRA activity (*Aeromonas*) (27, 41-43). The genomic analyses of these phylogenetically diverse DNRA-catalyzing isolates confirmed that the possession of *nrfA* or *nirB* is necessary for DNRA phenotype (32, 34). All four isolates utilizing lactate as the electron donor were *nrfA* genotype. *Enterobacter* sp. DNRA5 and *Klebsiella* sp. DNRA6 lacking *nrfA* failed to grow on lactate at NO_2_^−^-reducing condition, suggesting that NrfA-type nitrite reductase is needed for respiratory NO_2_^−^ reduction to NH_4_^+^. Thus, physiological functions of NirB-catalyzed NO_2_^−^-to-NH_4_^+^ reduction in these organisms may be NAD^+^ regeneration for fermentation, detoxification of NO_2_^−^, and/or assimilatory reduction, as previously suggested (14, 32). Neither *nirS* nor *nirK* was found in any of the sequenced draft genomes; however, a *nosZ* gene was recovered in the genomes of the *nrfA-* possessing *Bacillus* sp. DNRA2. Incubations on NO_3_^−^ reduction with and without C_2_H_2_ confirmed N_2_O reduction activity in this isolate amid active DNRA, which was probably catalyzed by the NosZ encoded by this gene.

The most easily recognizable common phenotype of the DNRA isolates was NO_2_^−^ accumulation before NH_4_^+^ production suggesting NO_3_^−^ repression of NO_2_^−^-to-NH_4_^+^ reduction. This phenotype has been consistently observed in previously studied DNRA bacteria (34, 35, 44). These previous studies attributed the NO_3_^−^-induced repression to transcriptional regulation involving NO_3_^−^ sensor proteins NarQ and NarX, and the transcript abundances of *nrfA* in *Escherichia coli*, and both *nrfA* and *nirB* in *B. vireti* were significantly lower when supplied with higher NO_3_^−^ concentrations. In agreement with these previous studies, *nrfA* in *Citrobacter* sp. DNRA3 and *nirB* in *Enterobacter* sp. DNRA5, exhibited at least 5.8-fold higher transcription after NO_3_^−^ depletion than before (*p*<0.05). Further, the results from the resting-cell experiments with these isolates showed clear indications that the presence of NO_3_^−^ at sub-millimolar concentration was sufficient to inhibit activities of expressed NrfA-type nitrite reductase. Whether the apparent inhibition was due to the redirection of electron flow analogous to what was observed with NosZ-catalyzed N_2_O reduction in presence of O2 or inhibition of NrfA enzyme itself, however, cannot be determined and is outside the scope of the current study (45). In denitrifiers, such NO_3_^−^-mediated repression of dissimilatory NO_2_^−^ reduction, either via transcription regulation or enzyme inhibition, has not yet been reported, and near-stoichiometric NO_2_^−^ accumulation during NO_3_^−^ reduction has been observed only as isolated cases (46-48). Therefore, as long as NO_3_^−^ is present in soil matrices harboring diverse denitrifiers and DNRA-catalyzing organisms, NO_2_^−^ produced from NO_3_^−^ would be reduced mostly to N_2_O and N_2_ via denitrification, with DNRA phenotype remaining silent.

The environmental physicochemical parameter that has been most frequently associated with DNRA activity is the C-to-N ratio. Multiple experimental evidences from culture-based experiments and field measurements have supported that DNRA is favored at high C-to-N ratios, i.e., electron acceptor limiting condition, while denitrification is favored at low C-to-N ratios, i.e., electron donor limiting condition (17, 19, 22). The observations from the incubation of the two isolates at the two different C-to-N ratios indicated that the NO_3_^−^ repression of NO_2_^−^-NH_4_^+^ reduction activity may actually be directly linked to this C-to-N ratio regulation of DNRA activity in the environment. The C-to-N ratio of soils or sediments are often inversely related to the NO_3_^−^ contents (49). In the soils with low C-to-N ratios, the NO_2_^−^-to-NH_4_^+^ reduction may thus be deactivated in the DNRA-catalyzing organisms due to the high NO_3_^−^ contents while NO_2_^−^-to-N_2_O/N_2_ reduction activity remains intact in denitrifiers cohabiting the ecological niches. Even with abundant DNRA-catalyzing population, N_2_O and N_2_ would still be the dominant terminal product of NO_3_^−^ reduction in such soils. Thus, what was previously regarded as the effect of the C-to-N ratio on the denification-vs-DNRA competition may be, at least in part, explained as the consequence of NO_3_^−^ inhibition of *sensu stricto* DNRA (39).

Production of N_2_O has been consistently observed in non-denitrifying organisms with DNRA phenotypes, with recovery of up to ~50% of NO_3_^−^-N as N_2_O-N (28, 32, 44, 50). Likewise, all DNRA-catalyzing isolates examined in this study produced N_2_O during the course of NO_3_^−^ reduction to NH_4_^+^ despite the absence of *nirS* or *nirK* gene in their genomes. Hypotheses surrounding the mechanisms of N_2_O production from DNRA have invariably involved NO (32, 35, 44). Considering that *norB, norV*, and/or *hmp* genes were identified in the genomes of DNRA isolates, it is plausible to regard NO as the precursor of N_2_O in these organisms; however, the mechanism leading to NO production from NO_3_^−^ or NO_2_^−^ remains unclear. The absence of N_2_O production before NO_3_^−^ depletion in all six isolates suggested against involvement of NapA- or NarG-type nitrate reductases. The more plausible source of NO would be NO_2_^−^-to-NH_4_^+^ reduction by NrfA- and NirB-type nitrite reductases, although reaction mechanism leading to formation of NO as a byproduct has not been elucidated for either enzyme (44, 51). The results from the chemostat experiments with *Enterobacter* sp. DNRA5 further support this hypothesis, as detectable N_2_O production was observed only at the high C-to-N ratio operating condition where active NO_2_^−^-to-NH_4_^+^ reduction occurred.

Another noteworthy observation regarding N_2_O production by the DNRA-catalyzing organisms was the absence of detectable N_2_O production in the NH_4_^+^-producing *Citrobacter* sp. DNRA3 chemostat (high C-to-N ratio), which suggested that the NO_3_^−^ or NO_2_^−^ concentrations in the surrounding environment may have positive correlation with N_2_O formation by *nrfA*-possessing organisms. In line with this observation, elevated N_2_O production upon incubations with media with lower C:N ratios, i.e., higher NO_3_^−^ concentration, was previously observed in *nrfA*-utilizing isolates of *Citrobacter* and *Bacillus* genera (28). Together with the finding that *Bacillus* sp. DNRA2 actively consumed N_2_O simultaneously with NO_2_^−^-to-NH_4_^+^ reduction, this observation suggests that N_2_O produced from NrfA-mediated DNRA may be negligible in complex soil microbiomes.

## MATERIALS AND METHODS

### Soil sampling and initial characterization

The agricultural soil used in this study was sampled from an experimental rice paddy field located at Chungnam National University (CNU) agricultural research site in Daejeon, Korea (36°22’01.6”N 127°21’14.3”E) in October 2018. Harvesting had been complete and there was no standing water at the time of sampling. Cover soil and plant materials were carefully removed before sampling and approximately 1 kg of soil at 5 to 30 cm depth from the surface was collected with a stainless tubular soil sampler with an inner diameter of 2 cm. The sampled soils were transported to the laboratory in coolers filled with ice and stored at 4°C until use. The physico-chemical characteristics of this soil, including the pH, textures, and total carbon and nitrogen contents were analyzed using standardized protocols.

### Culture medium and growth conditions

The minimal salts medium (MSM) for enrichment, isolation, and cultivation of soil DNRA bacteria was prepared by adding, per L of deionized distilled water, 10 mmoles NaCl, 3.24 mmoles Na_2_HPO_4_, 1.76 mmoles KH_2_PO_4_, 0.1 mmoles NH_4_Cl, and 1 mL of 1000x trace element stock solution (52). For enrichment and isolation, R2A broth (Kisanbio, Seoul, Korea) was added to the medium as growth supplements. To minimize the interference of NH_4_^+^ derived from mineralization of organic nitrogen in the ensuing DNRA-screening process, R2A broth concentration in the medium was limited to 6.2 mg L^−1^. The pH of the medium was adjusted to 7.0. Pure-culture incubation and experiments were performed with 100 mL medium dispensed to 160-mL serum vials. The serum bottles were flushed with N_2_ gas (≥99.999%, Special Gas Inc., Daejeon, Korea) for 10 min and sealed with black butyl-rubber stoppers (Geo-Microbial Technologies, Inc., Ochelata, OK) and crimped with aluminum crimp seals before autoclaving. The degassed filter-sterilized 200X vitamin stock solution was then added to the medium (52). Immediately before inoculation, sodium lactate or glucose was added as the electron donor and organic carbon source and KNO_3_ as the electron acceptor. Lactate and glucose were chosen as the non-fermentable and fermentable electron donors, respectively, as both substrates had been previously reported to support DNRA reactions (17, 53, 54). Enrichment with acetate as the electron donor was also attempted, but failed to yield any DNRA-positive isolates. Thus, acetate was not further considered as a potential electron donor in this study. The culture bottles were incubated with shaking at 150 rpm in dark at room temperature (25°C). Agar plates were prepared by adding 15 g L^−1^ Bacto^™^ Agar (Becton Dickinson, Franklin Lakes, NJ) to the liquid medium prepared with elevated concentrations of KNO_3_ (12 mM) and sodium lactate or glucose (120 mM as C). The 96-well plates for DNRA-screening were prepared by distributing 200-μL aliquots of the prepared culture medium to the wells in a UV-sterilized 96-well clear flat bottom microplates (Corning Inc., Corning, NY). Agar plate and 96-well plate were prepared and incubated in an anaerobic chamber (Coy Laboratory Products, Inc., Grass Lake, MI) with atmosphere consisting of 96% N_2_ and 4% H_2_.

### High-throughput DNRA phenotype screening

A simple novel high-throughput screening method was developed in this study for isolating DNRA-catalyzing organisms from agricultural soils without apparent DNRA activity (Fig. 6). Anoxic soil enrichments were prepared in 250-mL Erlenmeyer flasks (Duran Group, Wertheim, Germany) in the anaerobic chamber. The rice paddy soil was suspended at 1:10 w/v soil-to-medium ratio in 200 mL MSM amended with 2 mM KNO_3_ and 6.67 mM sodium lactate or 3.33 mM glucose (20 mM total C concentration) and incubated for two weeks in dark without shaking. The aqueous NO_3_^−^-N, NO_2_^−^-N, and NH_4_^+^-N concentrations were measured to confirm depletion of the electron acceptors and to check the extent of DNRA reaction in the enrichments. Serial dilutions of the soil enrichment cultures were spread onto agar plates, and after incubation in the anaerobic chamber, single colonies were picked to inoculate the 96-well plates loaded with fresh medium containing 1 mM NO_3_^−^ and 3.34 mM lactate or 1.67 mM glucose. The inoculated 96-well plates were covered with an optical adhesive cover (Applied Biosystems, Foster City, CA) to prevent contamination and evaporation of the culture medium and incubated for a week.

**Fig. 6.**
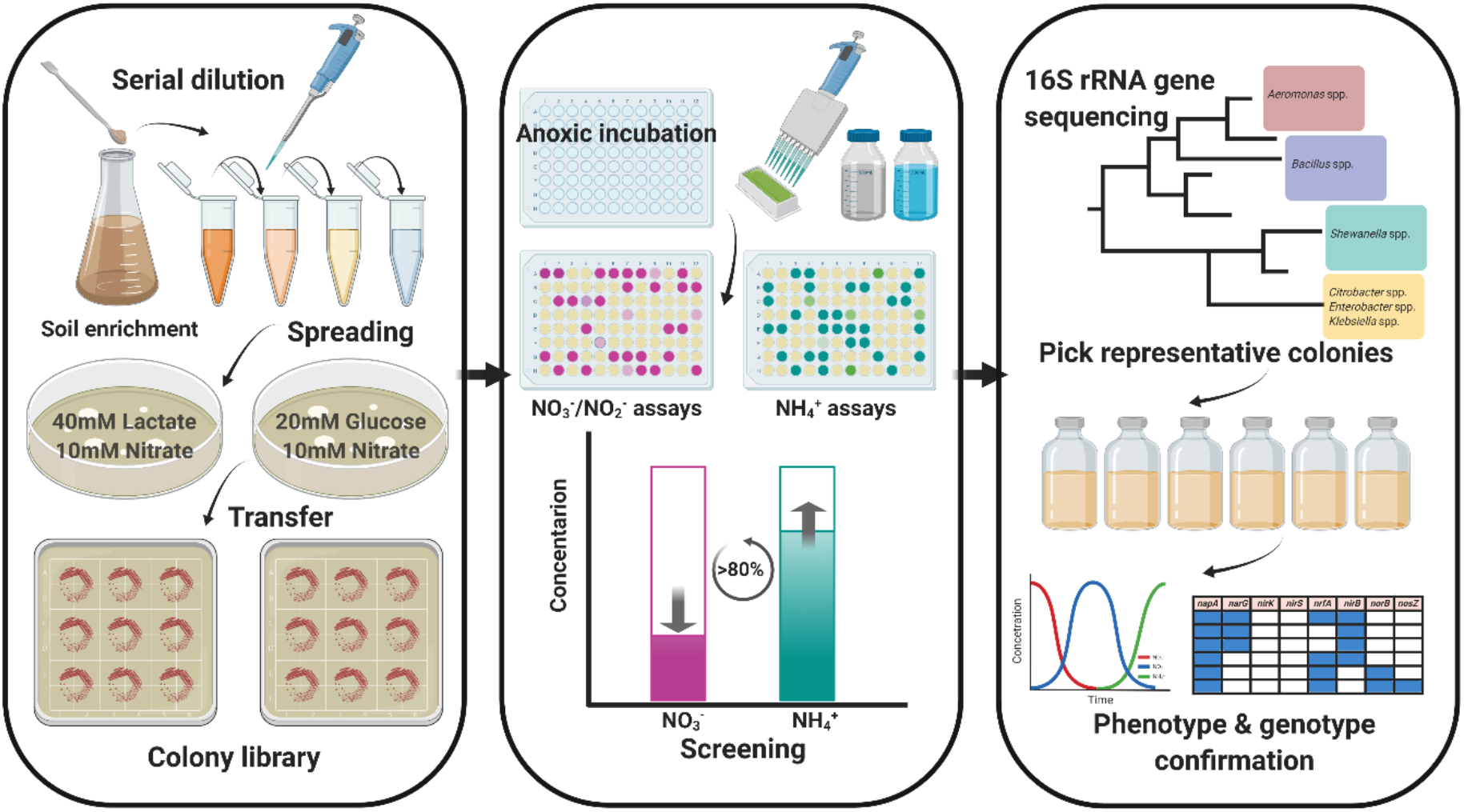
The schematic overview of the high-throughput screening methods developed for isolation of DNRA-catalyzing organisms from agricultural soil

After incubation, 100 μL of the 200-μL culture in each well was transferred to its corresponding position on a new 96-well plate. The absorbances at 600 nm, 660 nm, and 540 nm were determined using a Sunrise microplate reader (Tecan, Männedorf, Switzerland) with one of the duplicated plates to measure the possible interference of cell turbidity to the colorimetric determination of inorganic nitrogen concentrations. This plate was used for screening of the wells with increased NH_4_^+^ concentrations indicative of DNRA activity, using salicylate-nitroprusside chemistry. To each well, 80 μL of the color reagent (containing 0.2 M sodium hydroxide, 1 M sodium salicylate and 5.88 mM sodium nitroprusside dihydrate) and 20 μL of 5.1 mM sodium dichloroisocyanurate solution were sequentially added. The absorbance at 660 nm was measured after 30-minute incubation at 25°C, and the wells with OD_660nm_ values higher than 1.6 (equivalent to 0.8 mM NH_4_^+^) after subtracting OD_660nm_ resulting from cell turbidity were considered as positives. The duplicated 96-well plate was used for sequential measurements of NO_2_^−^-N and NO_3_^−^-N (55). The Griess reagent was added to each well to a total volume of 200-μL and the absorbance at 540 nm was measured after 30 min of incubation at 25°C for determination of NO_2_^−^-N concentration. The NO_3_^−^-N concentration was determined after the initial Griess assay. After reducing NO_3_^−^ to NO_2_^−^ by adding per well 20 μL of 1% w/v vanadium(III) chloride (VCl3; Sigma-Aldrich) prepared in 1 M HCl aqueous solution, the absorbance at 540 nm was measured to obtain the NO_3_^−^-plus-NO_2_^−^ concentration, from which the NO_2_^−^ concentration was subtracted. The colonies corresponding to the wells with NH_4_^+^ concentrations higher than 0.8 mM and absence of NO_2_^−^ and NO_3_^−^ were transferred to gridded fresh agar plates and stored at 4°C until use.

### Characterization of DNRA isolates

The partial 16S rRNA genes of these candidate DNRA isolates were amplified with the 27F/1492R primer set and sequenced to identify their phylogenetic affiliations (GenBank accession number: MT426123-64). Based on these initial 16S rRNA sequencing data, one isolate per genus was selected and subjected to further analyses. The DNRA activity of each isolate was confirmed by incubating the isolate with NO_3_^−^ as the sole electron acceptor in 100 mL MSM in 160-mL serum bottle with and without 10% v/v C_2_H_2_ in the anoxic N_2_ headspace. C_2_H_2_ inhibits N_2_O reduction by NosZ, thus enabling observation of N_2_O production from NO: and/or NO_2_^−^ reduction in a NosZ-harboring organism (56). The changes to the dissolved concentrations of NO_3_^−^-N, NO_2_^−^-N, and NH_4_^+^-N, headspace concentrations of N_2_O, and the microbial growth (OD600nm) were monitored until no further change was observed.

After incubation, culture sample was collected from each of these six DNRA-positive isolates and the genomic DNA was extracted using the DNeasy^®^ Blood & Tissue Kit (Qiagen, Hilden, Germany) according to the manufacturer’s protocol. Genome sequencing was performed using the HiSeq4000 platform (Illumina, San Diego, CA) at Macrogen Inc. (Seoul, Korea). Quality trimming and removal of adapter sequences from raw reads was performed using Cutadapt v2.9 and *de novo* assembly using SPAdes (v3.14.0) with the minimum contig length set to 200 bp (57, 58). The quality of the draft genomes was assessed using the CheckM software v1.0.18 (59). The NCBI Prokaryotic Genome Annotation Pipeline (PGAP) was used for genome annotation (60). The presence or absence of the nitrogen dissimilation functional genes was double-checked by running *hmmsearch* command of the HMMER software package v3.1b1 with the hidden Markov models (HMM) downloaded from the FunGene database (http://fungene.cme.msu.edu/) accessed on 10/14/2019 (61). This process ensured that the missing genes were not due to incompleteness of the draft genomes. The genes encoding the regulatory proteins putatively involved in nitrogen dissimilation were also searched for in the annotated genome. The draft genome sequences of the six isolates were deposited to the NCBI’s GenBank database (accession numbers: JABAIU000000000, JABAIT000000000, JABAIS000000000, JABAIR000000000, JABAIQ000000000, and JABAIP000000000).

### NO_3_^−^ inhibition of NO_2_^−^-to-NH_4_^+^ reduction

*Citrobacter* sp. DNRA3 carrying single copies of *nrfA* and *nirB* genes and *Enterobacter* sp. DNRA5 carrying *nirB* genes were selected to further examine whether and how NO_3_^−^ affect NO_2_^−^-to-NH_4_^+^ reduction. The resting-cell NO_3_^−^ and NO_2_^−^ reduction activities were examined with the cells harvested from the two distinct phases of DNRA reaction, i.e., NO_3_^−^-to-NO_2_^−^ and NO_2_^−^-to-NH_4_^+^ reduction. The DNRA-catalyzing isolates were grown with 5 mM NO_3_^−^ as the electron acceptor and 40 mM lactate or 10 mM glucose as the electron donor and carbon source. The cells were harvested before and after NO_3_^−^ depletion. Cell pellets were collected by centrifuging 200-mL culture at 10,000Xg for 20 min at 4°C, and re-suspended in 10 mL MSM. One milliliter of the cell suspension was added to a 160-mL stopper-sealed serum vial containing 100-mL of fresh MSM with N_2_ headspace. Chloramphenicol (water-soluble; Sigma-Aldrich) was added to the final concentration of 25 μg mL^−1^ to arrest *de novo* protein synthesis (62). These cell suspensions were then amended with 2 mM NO_3_^−^ or NO_2_^−^ and 6.67 mM lactate (*Citrobacter* sp. DNRA3) or 3.34 mM glucose (*Enterobacter* sp. DNRA5). The rates of change in the amounts of NO_2_^−^ were measured and normalized with the protein mass of the resting-cell cultures.

To observe the effects of changing NO_3_^−^ and NO_2_^−^ concentrations on transcriptional expression of the nitrite reductase genes directly relevant to DNRA, transcript abundances of *nrfA* and *nirB* genes in *Citrobacter* sp. DNRA3 and *nirB* gene in *Enterobacter* sp. DNRA5 were monitored as the cells were grown with 1 mM NO_3_^−^ and 3.34 mM lactate or 1.67 mM glucose. Collection and treatment of the samples, including extraction, purification, and reverse transcription processes, were performed using established protocols (17). Quantitative polymerase chain reaction was performed with a QuantStudio 3 Real-Time PCR instrument (Thermo Fisher Scientific, Waltham, MA) using SYBR Green detection chemistry, targeting *nrfA* and *nirB* in *Citrobacter* sp. DNRA3 and *nirB* in *Enterobacter* sp. DNRA5, as described in detail in the supplemental material.

### Batch and chemostat incubation of the DNRA-catalyzing isolates with varying C-to-N ratios

Batch cultures of *Citrobacter* sp. DNRA3 and *Enterobacter* sp. DNRA5 were prepared with two different C (carbon in the organic electron donor)-to-N (nitrogen in NO_3_^−^) ratios, and DNRA reaction was observed in these vessels. For high-C-to-N incubation, the culture medium was initially prepared with 0.2 mM NO_3_^−^ and 5 mM lactate or 2.5 mM glucose, and after each NO_3_^−^/NO_2_^−^ depletion event, the culture vessels were amended with an additional batch of 0.2 mM NO_3_^−^. For low-C-to-N incubation, the initial medium contained 2 mM NO_3_^−^ and 0.2 mM lactate or 0.1 mM glucose, and the organic electron donors were replenished upon depletion, indicated by discontinued NO_3_^−^ reduction. The concentrations of NO_3_^−^, NO_2_^−^, and NH_4_^+^ were monitored throughout the incubation periods.

The chemostat cultures of the DNRA isolates were set up with 300-mL culture in continuously stirred 620-mL glass reactor fed fresh medium at a dilution rate of 0.05 h^−1^ (Fig. S5). The medium bottle and the reactor vessel were consistently purged with N_2_ gas to maintain anoxic culture conditions during incubation. The reactor was operated with high (10 mM lactate or 5 mM glucose and 2 mM NO_3_^−^ in the feed) and low (0.2 mM lactate or 0.1 mM glucose and 2 mM NO_3_^−^) C-to-N ratios. The concentrations of NO_2_^−^, NO_3_^−^, and NH_4_^+^ in the effluent was monitored until the reactor reached steady state, as indicated by three statistically similar NO_2_^−^, NO_3_^−^ and NH_4_^+^ concentrations measured with 6-hour intervals. The N_2_O production rate was measured after steady state was established by closing the gas inlet and outlet of the reactor and monitoring linear N_2_O production.

### Analytical methods

The concentrations of NH_4_^+^, NO_2_^−^, and NO_3_^−^ were determined calorimetrically. Upon each sampling event, 1-mL sample was extracted with a disposable syringe and the cell-free supernatant was subjected to spectrophotometric assays. The NH_4_^+^-N concentration was measured using the salicylate method and the NO_2_^−^-N and/or NO_3_^−^-N concentrations were determined using the Griess method (33, 63). Headspace N_2_O concentrations were determined using a HP 6890 series gas chromatograph equipped with a HP-PLOT Q column and a ^63^Ni electron capture detector (Agilent Technologies, Santa Clara, CA)(64). Helium (≥99.999%, Special Gas Inc., Daejeon, Korea) and CH4 (5%)/ Ar (95%) mixed gas were used as the carrier gas and the make-up gas, respectively. The injector, oven and detector temperature were set to 200, 85, 250°C, respectively. Assuming equilibrium between the aqueous and gas phases, the total amounts of N_2_O-N in reaction vessels were calculated using the dimensionless Henry’s law constant of 1.68 at 25°C (65). The concentrations of glucose, lactate and acetate were measured using a Prominence high-performance liquid chromatograph (Shimadzu, Kyoto, Japan) equipped with an Aminex^®^ HPX-87H column (Bio-Rad Laboratories, Inc., Hercules, CA). Protein concentrations were determined with the Quick Start^™^ Bradford Protein Assay kit (Bio-Rad Laboratories, Hercules, CA) using a concentration series of bovine serum albumin solution as standards.

### Statistical analysis

With the exception of the chemostat experiments, all incubation experiments were performed in triplicates and the data presented as the averages of the triplicate samples along with their standard deviations. Statistical analyses were performed using the R software v.3.6.3 (www.r-project.org), where one-sample Student’s *t*-tests determined statistical significance of temporal changes in transcript copy numbers or N species concentrations. A *p*-value threshold of 0.05 was applied.

## Acknowledgement

This work was supported by the National Research Foundation of Korea (2020R1C1C1007970) and USDA-NIFA grant (2014-67019-21614). The authors were also financially supported by the Brain Korea 21 Plus Project (21A20132000003).

